# Discrimination of the hierarchical structure of cortical layers in 2-photon microscopy data by combined unsupervised and supervised machine learning

**DOI:** 10.1101/427955

**Authors:** Dong Li, Melissa Zavaglia, Guangyu Wang, Yi Hu, Hong Xie, Rene Werner, Ji-Song Guan, Claus C. Hilgetag

**Author notes:** Corresponding author: Prof. Claus C. Hilgetag, PhD, Professor & Director, Institute of Computational Neuroscience, University Medical Center Eppendorf, Martinistraße 52, 20251 Hamburg, Germany, Website: www.uke.de/icns. These authors contributed equally to this work (Dong Li, Melissa Zavaglia, Guangyu Wang).

## Abstract

The laminar organization of the cerebral cortex is a fundamental characteristic of the brain, with essential implications for cortical function. Due to the rapidly growing amount of high-resolution brain imaging data, a great demand arises for automated and flexible methods for discriminating the laminar texture of the cortex. Here, we propose a combined approach of unsupervised and supervised machine learning to discriminate the hierarchical cortical laminar organization in high-resolution 2-photon microscopic neural image data without observer bias, that is, without the prerequisite of manually labeled training data. For local cortical foci, we modify an unsupervised clustering approach to identify and represent the laminar cortical structure. Subsequently, supervised machine learning is applied to transfer the resulting layer labels across different locations and image data, to ensure the existence of a consistent layer label system. By using neurobiologically meaningful features, the discrimination results are shown to be consistent with the layer classification of the classical Brodmann scheme, and provide additional insight into the structure of the cerebral cortex and its hierarchical organization. Thus, our work paves a new way for studying the anatomical organization of the cerebral cortex, and potentially its functional organization.

## Introduction

Although the mammalian cerebral cortex is not particularly thick, its laminar structure is remarkably complex^1-5^. More than a century ago, the classical Brodmann scheme identified six cortical layers, followed more recently by findings of sub-layers within them^6,7^, indicating the existence of a hierarchical (encapsulated) organization.

A hierarchical organization exists on many levels in the brain, for instance with regard to the anatomical connectivity of cortical areas^8,9^, their functional connectivity^10^, or intrinsic time scales across primate cortex^11^. The motivation to study such a hierarchical neural organization ultimately derives from the desire to understand brain function. With respect to the cerebral cortex, a number of studies have shown functionally discriminable properties of cortical layers. For instance, in rat primary somatosensory cortex, the response latency to external stimuli in layer Va is significantly different from layers Vb, IV and VI^12^; and in mouse cortex, memory trace neurons are found primarily in layer II/III rather than layer IV^13^. From a technical point of view, calcium imaging or gene expression imaging in brain tissue achieves a spatial resolution in the micrometer range^14-23^, allowing detailed analysis of, for instance, the mouse cortex (thickness around 1 mm^24^). The use of modern imaging techniques for further investigation of the laminar hierarchy of cerebral cortex, therefore, promises further insights into brain function, but corresponding analysis approaches have to be able to deal with, and take advantage of, currently recorded massive high-resolution neural imaging data.

The conventional way of manually labelling cortical layers according to a classical atlas clearly faces problems in this era of massive high-resolution neuroimaging data^25^. Cortical laminar patterns vary from area to area, from species to species, and even from individual to individual within the same species. Whereas the increasing availability of a large amount of data is potentially a great advantage for research, it is too labor and time intensive to label the cortical layers for every location in every animal and research project. Moreover, manual labelling entails an observer-specific bias, particularly in areas without clearly visible traits for differentiating the layers, or when going beyond classically established layers to identify potential sub-layers. In addition, although the classical definition of laminar layers and their hierarchy contributed much to cortical research, by grouping neurons for comparable data analysis and computational modeling, it sometimes appears overly restrictive. For instance, it suggests that, in terms of hierarchical organization, Brodmann’s six layers can be considered to reside on the same hierarchy level, and further sub-layers on the levels below. However, particularly with respect to the cytoarchitecture of rodents, it has been found that Brodmann’s layers II and III display high anatomical similarity^25^, whereas Brodmann’s layer Va significantly differs from other sub-layers of layer V^1,26-30^. Thus, there is a great demand for neuroimaging data processing solutions that can address these challenges, by providing an automatic and objective differentiation, but also flexible definition of cortical laminar hierarchies.

Machine learning is a very promising approach in this regard, and existing findings of studies on conventional layer labelling and laminar properties of cortical neurons, such as morphological^31^, physical^32-36^, or functional^37^ properties of the neurons, already provide a range of well-characterized features that can be employed for machine learning-based laminar pattern discrimination.

Some efforts have already been made to automatically and objectively discriminate neurons belonging to different layers by supervised machine learning approaches^38-41^, yielding appealing laminar classifications; yet, the training data sets and methods are still limited by the framework of conventional atlases. The existing methods can, therefore, address the issue of labor intensity, but do not yet contribute to the other challenges. In contrast, unsupervised learning and clustering algorithms^42^ allow, in principle, for more flexibility. So far, however, existing work in this context still requires the predefinition of a desired number of layers and this aspect, in turn, limits the required layer and sub-layer definition flexibility. In the present work, we, therefore, propose combining unsupervised and supervised machine learning to discriminate the hierarchical organization of cortical layers. To illustrate the feasibility of the approach, the developed methodology is applied to 2-photon microscopic image data from mouse cortex. For small cortical foci, we use unsupervised clustering to identify the laminar cortical structure. During clustering, we do not demand a particular number of layers per hierarchical level, but automatically set up the level-specific layer structure. Supervised machine learning is then used to bridge the hierarchical clustering results obtained at different cortical locations and animals, that is, to ensure existence of a consistent layer labelling system across cortical locations. Manual labelling is, therefore, not required for the discrimination process. We, nevertheless, compare the classification results with existing laminar cortical schemes as part of an extensive evaluation to demonstrate plausibility of our layer discrimination results. In particular, using neuro-biologically meaningful features, we illustrate that our discrimination results are consistent with the classical Brodmann scheme of cortical layers, and in addition provide useful information on the hierarchical laminar organization in different cortical areas.

## Materials and Methods

### Imaging data description

#### Data acquisition

The used data was high-resolution 2-photon imaging mouse data previously described by Xie et al^13^. Specifically, the mouse strain is BAC-EGR-1-EGFP (Tg(Egr1-EGFP)GO90Gsat/Mmucd from the Gensat project, distributed from Jackson Laboratories. Animal care was in accordance with the institutional guidelines of Tsinghua University. 3-5 months old mice received cranial window implantation as previously described^13^; recording began one month later. To implant the cranial window, the animal was immobilized in custom-built stage-mounted ear bars and a nosepiece, similar to a stereotaxic apparatus. A 1.5 cm incision was made between the ears, and the scalp was reflected to expose the skull. One circular craniotomy (6-7 mm diameter) was made using a high-speed drill and a dissecting microscope for gross visualization. A glass-made coverslip was attached to the skull. For surgeries and observations, mice were anesthetized with 1.5% isoflurane. EGFP fluorescent intensity (FI) was imaged with an Olympus Fluoview 1200MPE with pre-chirp optics and a fast AOM mounted on an Olympus BX61WI upright microscope, coupled with a 2 mm working distance, 25x water immersion lens (numerical aperture, 1.05).

#### Data characteristics

For each mouse, usually 10 to 20 cortical locations were monitored. Each location was 510 μm × 510 μm broad and scanned until a depth of between 300 μm and 800 μm (see, for example, in Fig. 1), containing a few to over ten thousand neurons. The image resolution in *z* direction was always 2 μm; in-plane resolution was 0.996 μm × 0.996 μm. Each mouse was scanned once per day, for between a few and up to tens of days. In this study, we randomly chose sixty locations from eight mice, including 10 in motor areas, 7 in Posterior parietal (PTLp) areas, 18 in retrosplenial cortex (RSC) areas, 7 in primary somatosensory (SSp) areas, 6 in anterior medial visual (VISam) areas, 8 in primary visual (VISp) areas, 2 in unspecified visual (VIS) areas, 1 location on the boundary between Motor and SSp and 1 location around the junction of PTLp, SSp and RSC. Each location was named in such a way that it was distinguishable. Neuron positions in the images were automatically detected with high accuracy using the neuron network model described by Xie et al^13^.

**Figure 1.**
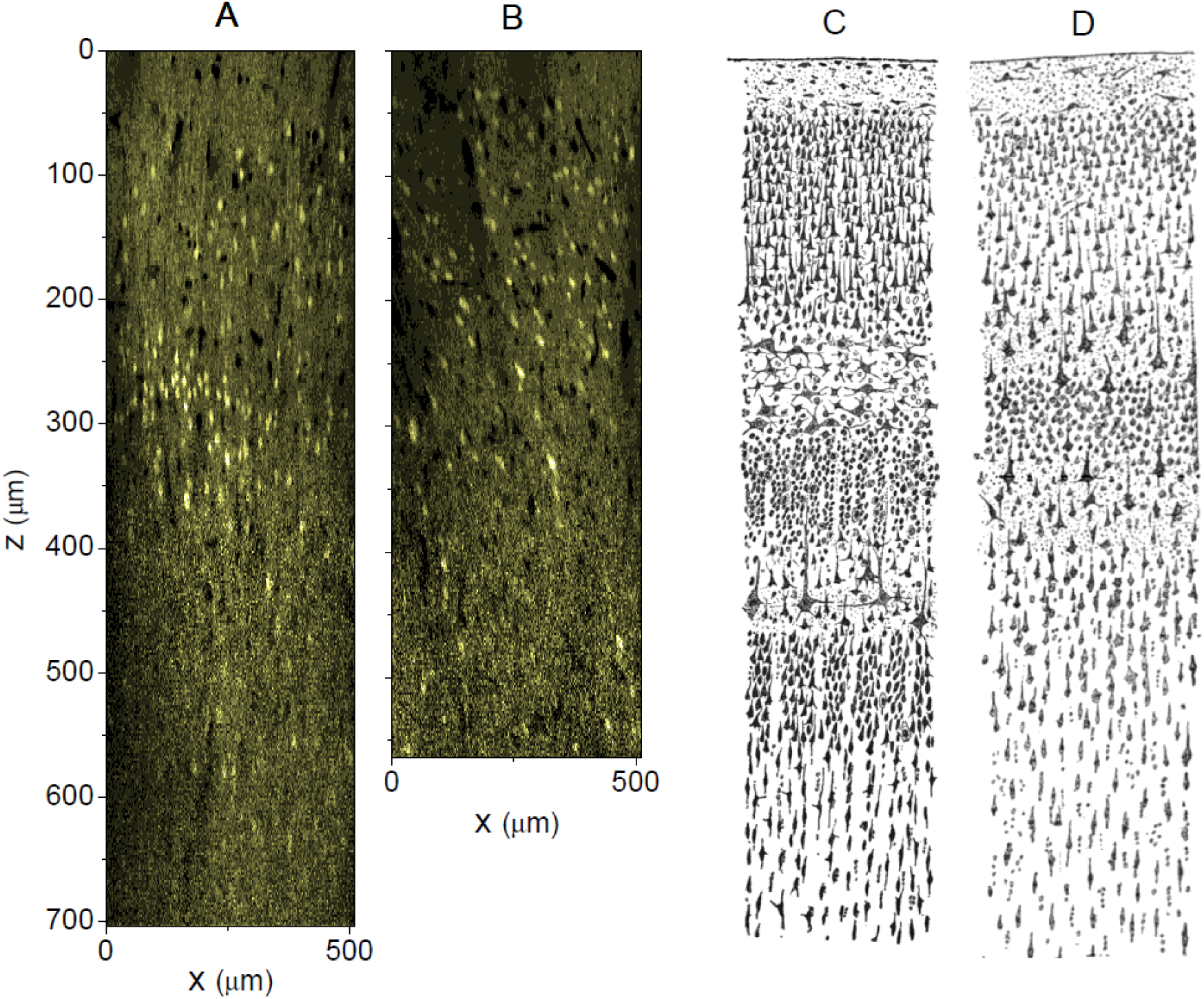
Example slices of x-z planes of (A) VISp area and (B) motor area of mouse brain in our study. For comparison purposes, we adapt the drawings of cortical lamination by Santiago Ramon y Cajal of (C) Nissl-stained visual cortex and (D) Nissl-stained motor cortex of a human adult^50^.

#### Manual labelling of Brodmann layers

For all sixty cortical locations, an experimental expert (G.W.) labelled those Brodmann layers (B-Layers) that could be unambiguously discriminated by eye in the location-specific image data. The labels were assigned to entire x-y image slices. Depending on the particular locations, the assigned Brodmann layers included (1) the superficial layer, (2) Brodmann layers II/III, (3) Brodmann layer IV, (4) Brodmann layer Va, (5) Brodmann layer Vb, (6) Brodmann layer VI, and (7) deeper parts. Each location was labelled with between four and seven of the Brodmann layers, depending on the area type and imaging specifics. For instance, some areas did not contain a visible layer IV or layer Va, and some image data sets did not cover all layers due to limited scanning depth. The manually assigned Brodmann layers were used for evaluation purposes of the automatic layer discrimination results. According to our experience, manual labelling uncertainty was approximately ±5 slices.

### Combined super- and unsupervised layer discrimination: principle

The overall pipeline of the proposed approach to discriminate the hierarchical laminar structure of the acquired data was shown in Fig. 2. The input observations were 3-dimensional image stacks *I:* (*x,y,z*) ∈Ω⊂ℤ^3^ ↦ *I*(*x,y,z*) with *I*(*x,y,z*) as image intensity at pixel location (*x,y,z*). Further, *I_z_* denoted the image slice at depth z and 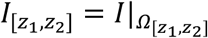 the restriction of *I* to 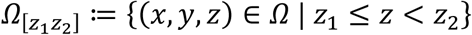, that was, all slices between z_1_ and z_2_. The overall aim was to discriminate layers 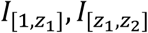, etc. and to represent their hierarchical organization in an objective and meaningful manner for all studied cortical locations. To this end, we developed a machine learning-based approach inspired by the human’s recognition process of cortical layers, consisting of two main parts. Within each cortical location, the discrimination of the laminar hierarchical structure was first obtained via clustering (i.e., unsupervised machine learning). The clustering results for the individual locations were then combined using supervised learning in order to transfer layer labels on particular hierarchical levels that were obtained for specific reference locations to the other locations. Thus, the supervised learning step ensured consistent layer labelling across different locations.

**Figure 2.**
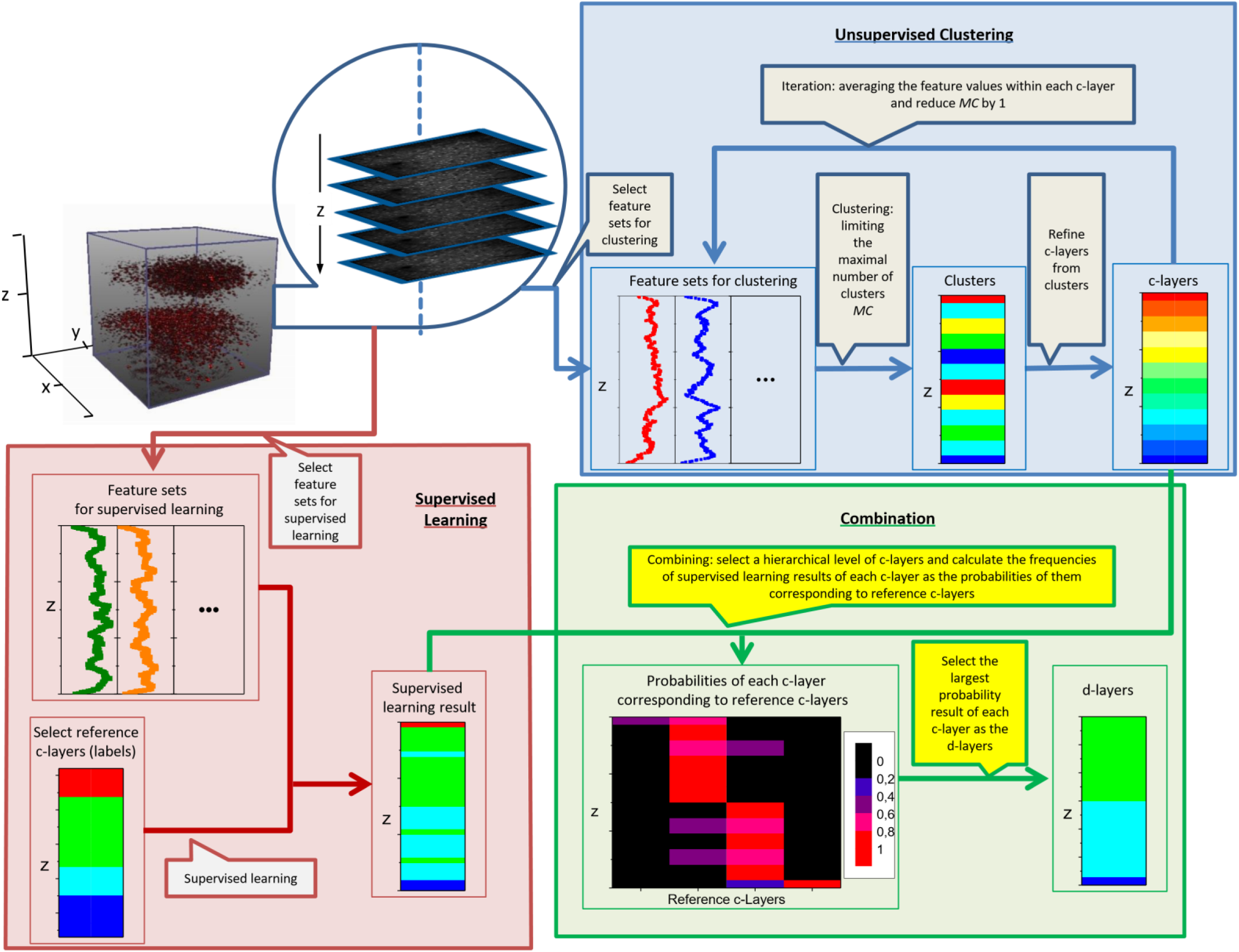
Proposed pipeline for the discrimination of hierarchical laminar cortical organization. The top-right panel shows the unsupervised clustering part, whereas the bottom-left panel shows the supervised learning part. The bottom-right panel shows how to combine the unsupervised and supervised learning results.

#### Terminology

The terminology *layer* referred to several but different concepts in this study. On an abstract level, layer simply meant a collection of z-slices 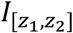. In this work, the layers that were manually labelled according to Brodmann’s atlas were referred to as Brodmann layers (*B-layers*). During unsupervised hierarchical clustering, we further obtained a certain number of layers on each hierarchical level, which were called *c-layer* (*c* is short for clustering), followed by an Arabian number that indicated the vertical order from superficial to deep. When c-layers were used as training data in the supervised learning part, they would be denoted as *reference c-layers*. Those c-layers, to which the layer labels have to be transferred to, were denoted as *target c-layers*. The results of this final discrimination would be called *d-layers* (*d* for discriminated), followed by an Arabian number, indicating its corresponding position in the reference c-layer dataset.

#### Part 1: Hierarchical clustering for unsupervised layer definition

For each cortical location, an iterative procedure was applied to compute the hierarchical structure of the data. Briefly speaking, we iteratively performed feature-based clustering of the z-slices of *I*, refined the clustering results in order to define iteration-specific c-layers and successively decreased the maximum possible number of clusters, with the maximum possible number of clusters subsequently denoted as *MC* (see the unsupervised learning panel in Fig. 2). The entire iteration process resulted in a global hierarchical tree, with the individual hierarchical levels representing the clustering results after the different iterations.

In detail, for each iteration, we first applied a classical hierarchical tree clustering. For the first iteration, *MC* was initialized by *MC_0_. MC_0_* can, in principle, be chosen arbitrarily, but not be too small. In our work, we chose *MC_0_*=10. Each clustering iteration then resulted in a certain number of clusters, with the clusters distributing along the z-direction of the image data set. Yet, depending on the considered features, different isolated c-layers might be assigned to the same cluster (e.g. layer 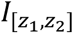 to cluster 1, 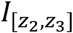 to cluster 2, and 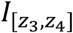 again to cluster 1 with *z*_1_*<z*_2_*<z*_3_*< z*_4_). As this was not the desired output, every isolated part of a cluster was defined as a specific c-layer, which meant that one cluster might be divided into more than one c-layer and the overall number of c-layers was larger or equal to the number of clusters *MC* (see again Fig. 2). After defining the c-layers for a specific iteration, the feature values for the individual z slices were replaced by the average values of the features corresponding to all the z-slices of the c-layer encompassing the considered z-slice. With these updated feature values and *MC* decreased by one, the next clustering iteration was initiated. This procedure was repeated until *MC* had a value of ‘one’. Thus, after each iteration, a series of new c-layers was computed that emerged from merging the c-layers of the previous iteration, i.e. the last lower hierarchical level.

From an implementation point of view, clustering was performed using the *linkage* algorithm of the MATLAB Statistics and Machine Learning Toolbox, with the *Euclidean* distance measure in feature space and the minimum variance algorithm (*ward*) as parameters.

#### Part 2: Across-location-consistent labelling of c-layers by supervised learning

To provide c-layer labels and a layer discrimination that was consistent for the different locations, we proposed combining the unsupervised clustering results with a supervised learning procedure (see the supervised learning panel and combination panel in Fig. 2). To this end, we first chose a reference dataset with rich laminar pattern, so as to cover the laminar characteristics and across-layer differences of as many layers as possible. Here, the reference data sets and respective reference c-layers were therefore selected from either VISp or SSp areas, that was, areas known to have well differentiable laminar layers^2,5^. The hierarchical levels on which reference and target c-layers were chosen for combination were in principle arbitrary, but the level should be similar for reference and target data set. In this study, for evaluation purposes, we always selected the 8^th^ hierarchical level for the reference location and the 7^th^ level in the target locations, as on those levels, the c-layer numbers were most similar to the number of expert-delineated B-layers.

To establish correspondence between the target and the reference c-layers, we first trained an ensemble of support vector machines (SVMs) based on the z-slices and slice-wise computed features of the reference data set. In total, *m* SVMs (in this work: *m =* 200 for each reference data set and each feature set) were trained, based on randomly selected slices of the reference data set and respective features.

The SVMs were then, individually for each target c-layer, applied to determine the corresponding reference c-layer. Let the reference c-layer labels be denoted by index *r =* 1,…, *r_max_* and the target c-layers by *t =* 1,…, *t_max_*. Each target c-layer *t* further consisted of *n_t_* image slices 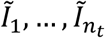 and respective features. Applied to each target c-layer slice, the SVM ensemble resulted in *m* slice-specific reference c-layer votes; repeating SVM application for all slices 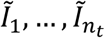 yielded in total *m · n_t_* votes. Let *n_r_* be the number of votes for reference c-layer *r*, the probability *p*(*r*) = *n_r_/*(*n_t_ · m*) was interpreted as similarity measure between target c-layer *t* and reference c-layer *r*. The reference c-layer with maximum similarity was then assigned to the target c-layer and referred to as d-layer.

The supervised machine learning part was also implemented in MATLAB, using the package LIBSVM^43^, using linear kernel SVMs with cost parameter = 1.

#### Definition of used features and feature sets

The following features were considered in this work and computed for the individual image slices *I_z_, z = 1*,…, *z_max_* of the 60 areas and corresponding image data sets described in section *Image data description*.

- **Neuron density:** Neuron density was the number of neurons per slice, normalized by the area of the slice (i.e., 510 μm × 510 μm). Neuron density was evaluated based on the automatic neuron detection described in the *Imaging data description* section.
- **Neuron size and neuron shape:** Precise computation and representation of neuron size and neuron shape required accurate segregation of the cell bodies, which is challenging by itself at the moment^44,45^ (see also, for instance, the 2018 Data Science Bowl challenge from the Kaggle competition^46^) and beyond the scope of this paper. Instead, we only used rough measures that reflect mean neuron size and 2D shape for the individual slices *I_z_*. As a result of the automated neuron detection, the locations of the neuron centres were known for the individual slices, and each neuron centre location was associated with a local maximum of the image intensity. To determine neuron size, the neuron soma boundary was estimated based on intensity, using half the image intensity value at the neuron centre relative to the background image intensity as a threshold; the sought size was defined as the number of the enclosed pixels, multiplied by the pixel spacing. To avoid negative influence of outlier values, the *robust fitting* function in MATLAB was used to finally calculate the average size of the neuron somata across all days of scanning for the particular location and image slice. To represent neuron shape, we computed a shape measure that reflected the difference of the shape of the 2D neuron soma area and a circle. Based on the soma delineation described above, the maximum soma diameter *d* was determined. The shape measure was then defined by (*πd^2^*)*/*(*4A*), with *A* denoting the soma size. Mean neuron shape measures were again computed as averages over all scanning days and neurons.
- **Neuron type:** Accurate discrimination of various types of neurons is currently also a topical challenge^40,47^ and, once again, not a direct target of this work. Instead, we only classified the detected neurons based on their shape and size (see definition above) into two groups, corresponding potentially to granule and pyramidal neurons. The classification was performed using a two-class Gaussian mixture model.
- **Image texture features**: To compare the performance of the neuron-related features for layer discrimination to other typical image-based features without direct relation to a neuro-biological meaning, we also computed and analysed the intensity co-occurrence matrices corresponding to the image slices. The following four image texture measures were considered (using the MATLAB function *graycoprops*): *Contrast* (average intensity difference between neighbouring pixels), *Correlation* (average correlation of intensity values of neighbouring pixels), *Energy* (also known as uniformity of an image), and *Homogeneity*.

#### Considered feature sets

In total, we defined six sets of features to demonstrate and compare their performance for layer discrimination: Feature set F1 (only neural density; i.e. one feature), F2 (neural density and mean neuron size; two features), F3 (neural density, the proportions of the two neuron classes, and the mean neuron sizes of the two classes; in total five features), F4 (F3, plus the mean size of all neurons; six features), F5 (neural density, mean neuron size, and the four image texture features; in total six features), and F6 (only the four image texture features).

### Evaluation strategies

In line with the structure of the proposed layer discrimination approach, the evaluation consisted of two parts: to evaluate the performed unsupervised clustering and to evaluate the supervised learning-based labelling of the target data c-layers. Methodologically, this leads to the tasks of measuring (P1) similarity between two computed laminar hierarchies, (P2) measuring similarity between a hierarchical clustering result and a (manually labelled) layer discrimination, and (P3) the direct comparison of two layer discrimination results, either manually or automatically obtained. The applied procedures are shown in Fig. 3 and detailed below.

**Figure 3.**
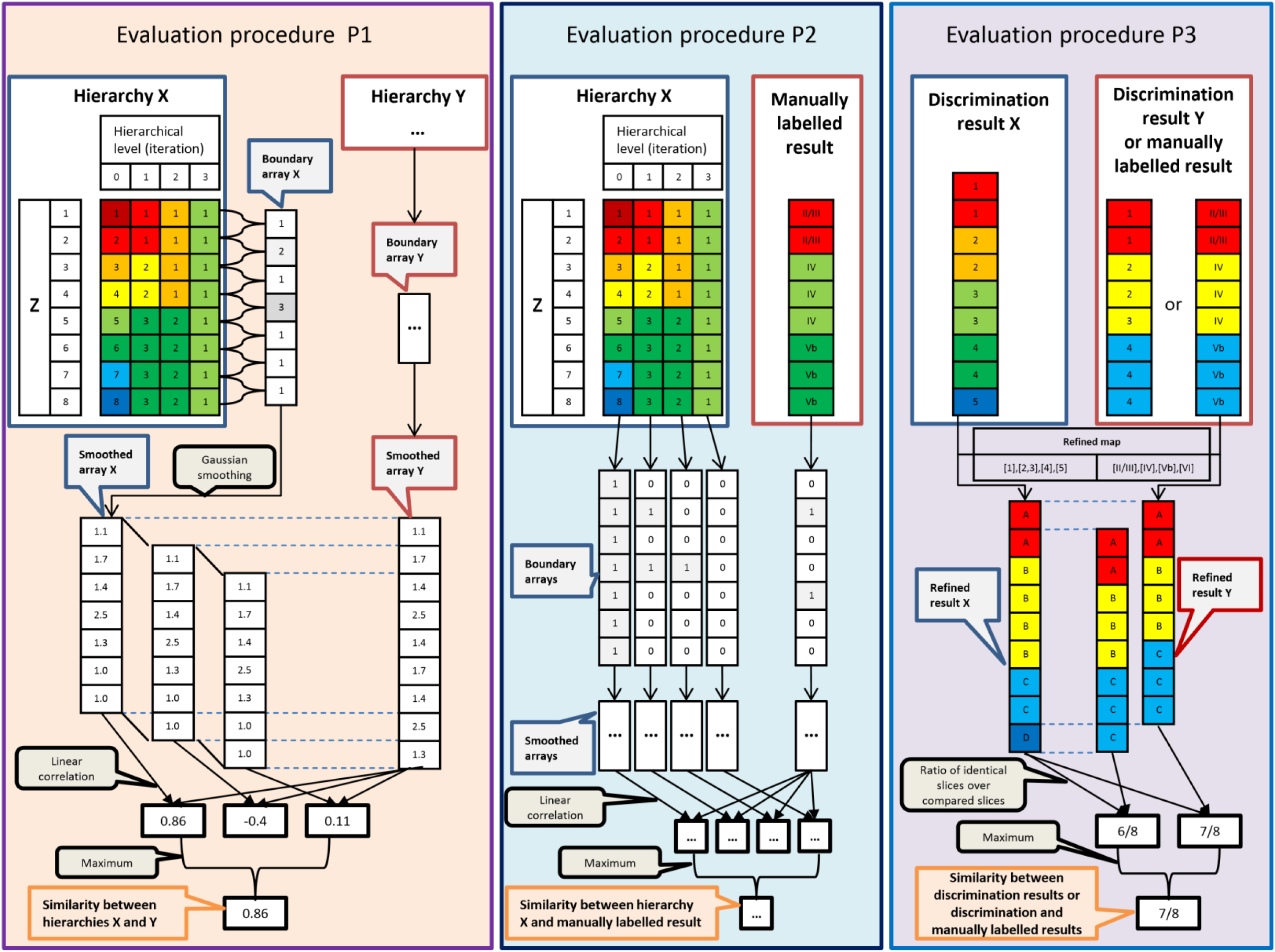
Illustration of applied evaluation procedures. The left panel shows the pipeline to evaluate the similarity between two hierarchical clustering results X and Y. X and Y could, for example, be two hierarchical trees obtained for one image data set but with different feature sets, or two hierarchy trees of data acquired at different locations. The middle panel illustrates the pipeline to evaluate similarity between a hierarchical clustering result and the corresponding manually labelled layers. The right panel shows the pipeline to evaluate two final discrimination results or a respective discrimination result and the corresponding manually assigned layer labels. Again, the two discrimination results can either indicate results computed for a single location but with different feature sets, or results from two different locations. For the latter case, the image data potentially contain a different number of slices.

#### Evaluation of unsupervised hierarchical clustering

To evaluate the results of the unsupervised hierarchical clustering, procedures (P1) and (P2) were used.

Procedure (P1) was applied to study the influence of the introduced feature sets on the hierarchy trees obtained by clustering. As shown in Fig. 3 (left panel), we first introduced a layer boundary function *ΔL^F^* that, for a given hierarchical tree after clustering based on a feature set *F*, assigns the particular tree level (traversing the tree in bottom-up direction) to each pair of consecutive slices (*I_z_, I_z+_*_1_) of the image data and area at hand, on which the slices for the first time were assigned to the same cluster. For example, in Hierarchy X in the left panel of Fig. 3, the first and the second slice merged at the first hierarchical level and, consequently, *ΔL*(*I_1_,I_2_*) = 1. The resulting layer boundary array

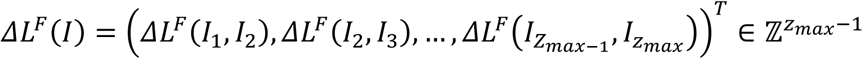

was Gaussian-smoothed (here: span of 5 elements). This smoothed boundary array 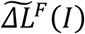 was the basis for subsequent evaluation steps. In particular, for each of the 60 areas, we evaluated the similarity κ of the boundary arrays 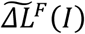 obtained for two different feature sets by calculating the linear correlation coefficient of the two boundary arrays.

Procedure (P2) was applied to measure the similarity between a hierarchical clustering result for a specific area and the manual layer labelling of the area. The manual labelling could be interpreted as a single-level hierarchy; thus, the boundary array *ΔL^man^*(*I*) given by the manual labelling only contained the elements 0 and 1 (see Fig. 3, middle panel). To, nevertheless, derive a similarity measure between *ΔL^man^*(*I*) and a feature set-specific hierarchical clustering result, we calculated a set of *binary* boundary arrays, one for each level of the hierarchical tree. If *l* denoted the hierarchical level considered, the corresponding binary boundary array 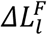 was defined by

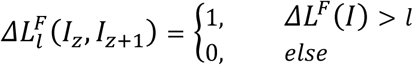

Similar to procedure (P1), after smoothing the binary boundary arrays, we calculated the linear correlation between 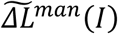 and the individual 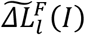 for all levels *l*. Finally, the maximum correlation coefficient Km of the coefficients obtained for the individual levels *l* was used to quantify the similarity between the laminar hierarchy and the manual labelling.

#### Evaluation of reference-based layer discrimination and label transfer

The technical procedure (P3) to evaluate the supervised learning-based transfer of layer labels from a reference to target data sets is described in the right panel of Fig. 3. Different to (P1) and (P2), we now assumed a specific layer discrimination and labelling result for an image data set *I* to be given that was to be compared to a ground truth layer discrimination of *I*. With *M^X^: ℤ → L_X_* mapping an image slice *I_z_* to a layer label 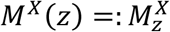 of an ordered label set *L_X_* that was specific to the discrimination result *X* and a similar function *M^Y^: ℤ → L_Y_* for the ground truth layer discrimination, the goal was to define a similarity measure between *M^X^* and *M^Y^*. As a first step, the potentially different label sets *L_X_* and *L_Y_* were mapped to a common label set *L_XY_*. For example, in the right panel of Fig. 3, *L_X_* represented the d-layer numbers 1, 2, etc, while *L_Y_* denoted Brodmann layers, indicated by Roman numerals. The different labels were mapped to the joint label set *L_XY_* and the respective labels A, B, etc. With 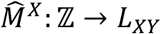 as *X*-specific mapping of *I_z_* into *L_XY_* (and analogously for the manually labelled ground truth *Y*), a similarity array was defined by

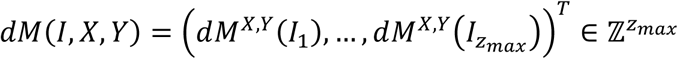

with

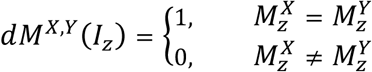

that served as the basis of the performed evaluation. In detail, the respective evaluation covered three aspects:

**Analysis of reference data and reference c-layers**: In a first step, we studied the properties and the quality of the reference data and reference c-layers, which served as the basis of the label transfer to different target data sets. As motivated above, the considered reference data were the 8 VISp and 7 SSp areas and the c-layers extracted after the 8^th^ iteration of unsupervised clustering using the F1 feature set. To alleviate interpretability of subsequent evaluation results (i.e. to ensure that d-layers assigned to target data sets could directly be compared to the target data set B-layers), we visually inspected the consistency of the reference c-layers and the manually assigned reference data B-layers. If necessary, reference c-layers were merged and apparent inconsistencies corrected. In Fig. 4F (data set M248L1), it could, for instance, be seen that c-layers 6 and 7 did not merge at the selected hierarchy level (the 8^th^ level), and both layers corresponded to the manually assigned layer 6 (i.e. B-layer VI); the two c-layers were therefore merged. The final mapping between the VISp reference c-layers and the manually assigned labels is summarized in Table 1; analogous information for SSp can be found in Table S1.

**Figure 4.**
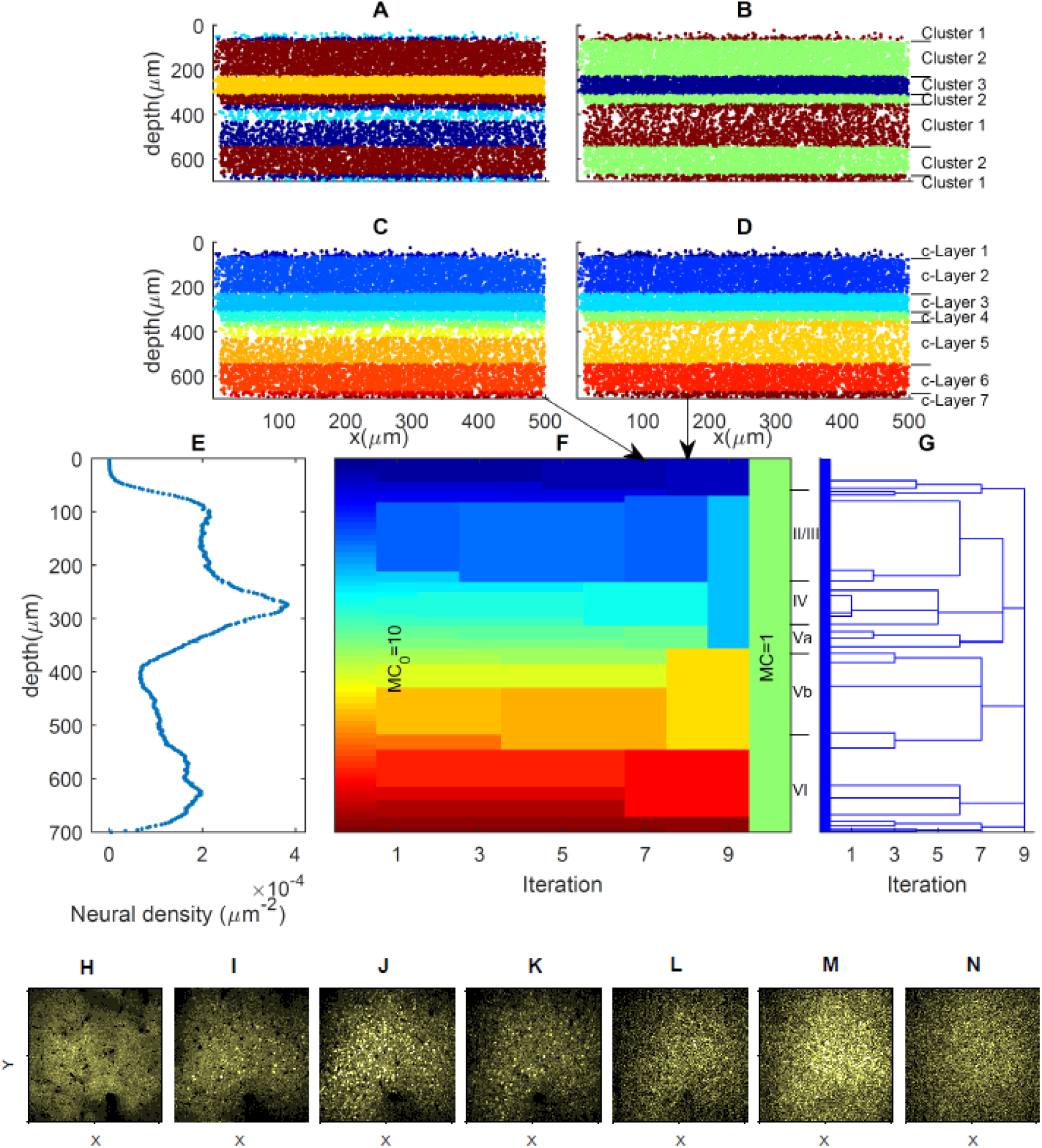
Example of unsupervised clustering (VISp area M248L1). Panels A and B show the distribution of the clusters as direct output of the original clustering algorithm and panels C and D the respective layers after 7^th^ and 8^th^ iterations. Panels F and G represent the whole iteration process and the hierarchical tree obtained with neural density as unique feature (panel E). The manually labelled layers are indicated to the right of panel F. The bottom panels show seven slice examples at z=40 μm (H), z=160μm (I), z=280μm (J), z=320μm (K) z=460μm (L), z=620μm (M), and z=690μm (N), corresponding to the seven c-layers on the eighth hierarchical level in panel F.

**Table 1:**
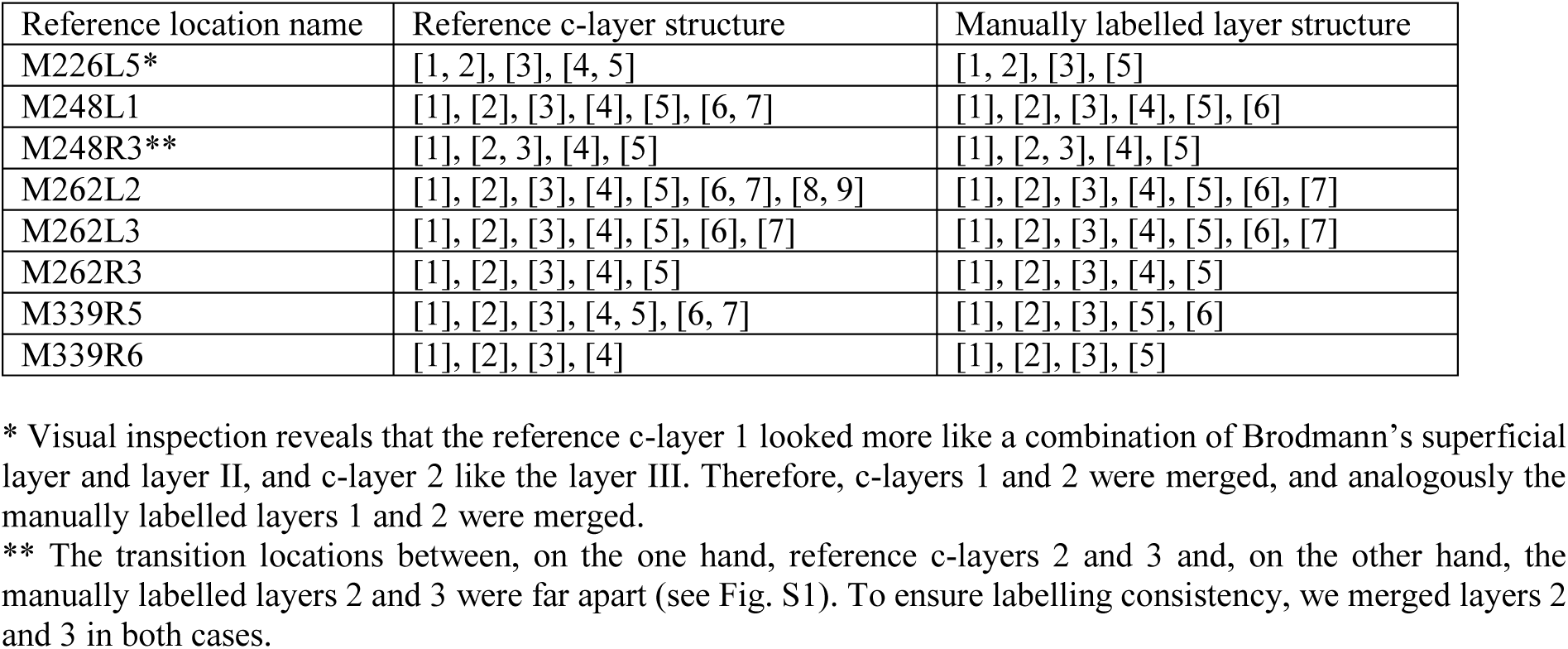
Refined maps between Reference c-layers from VISp areas (F1 feature set and the 8^th^ level on the hierarchy) and the manually labelled layers. Layer numbers grouped in parentheses were layers that were considered as one single layer during evaluation; see respective explanations for details.

Based on the refined reference c-layers and the refined B-layers labels, two quality measures, *n_B_* and *f_c_*_=B_, were introduced and evaluated for the individual reference data sets. *n_B_* denoted the total number of the manually labelled layers that the data set covered, which should be close to seven. Yet, due to limited scanning depth, the reference data sets did not necessarily cover all possible layers (see section *Image data description*), resulting in a smaller number, and limiting its usability as a reference data set. *f_c=B_* represented the number of slices where both the assigned refined c-layer and the assigned B-layer pointed to the same cluster, divided by the total number of slices of the data set. Referring to the aforementioned definitions, *f_c=B_* equaled the averaged value of the entries of the similarity array *dM*(*I, c, B*); here, *c* indicated reference c-layer-based and *B* the B-layer-based discrimination of the data set *I*. Thus, *f_c=B_* reflected the performance of the unsupervised clustering of *I*, with *f_c=B_ = 1* as ideal performance.

**Comparison of d- and B-layers (all locations):** To further evaluate the performance of supervised label transferring and the influence of the feature sets thereon, we transferred the refined reference c-layers to all locations and data sets. Thus, for each of the 15 reference data sets and 6 feature sets, 60 target datasets were considered and corresponding d-layers determined. This final d-layer definition was evaluated by slice-wise comparison of the d-layer- and the manually assigned B-layer-based discrimination, that was, by means of the measure *f_d=B_* that represented the fraction of the target data set slices with agreement of its associated d- and B-layers.

**Re-labelling of reference data:** As a third aspect, we constrained the aforementioned comparison to the reference datasets themselves serving as target datasets, but still using the different feature sets F1-F6. To measure the performance, we introduced the index 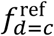 that represented the ratio of the number of reference dataset slices for which the original refined c-layers and the assigned d-layer were identical, and the total number of slices, i.e. the mean value of *dM*(*I, c, d*). Ideally, that was, if all slices within one reference c-layer had uniform but unique feature expressions, 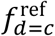 equals 1. Yet, this was not necessarily the case because, for instance, the unsupervised clustering approach comprised spatial connectivity constrains, while the supervised learning did not. The index 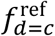 could, therefore, be understood as a measure to evaluate the relevance of individual features and appropriateness of the considered feature sets for supervised learning-based label transfer.

#### Analysis of cross-location similarity of discrimination results

As final part of the evaluation, we analysed the similarity of layer discrimination results between any pair of locations (i.e., in total 60 × 60 pairs) for unsupervised clustering as well as supervised learning and the different feature sets. From a neuro-biological perspective, image data from locations of the same area type could be expected to have more similar hierarchical laminar patterns than images from different area types; this should be reflected by automatic layer discrimination and appropriate features sets. For unsupervised clustering, we again calculated hierarchical laminar similarity using procedure (P1) and the linear correlation coefficient κ. For supervised learning, the d-layers were computed as described above. d-layer-based discrimination results computed for two (different) image data sets were then compared by means of the index 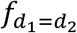 that resembled the introduced measure *f_d=B_* but using the d-layers of the first image (denoted by *d*_1_) and the d-layers of the second image (indicated by *d*_2_). Yet, the image data sets acquired at different locations contained in parts a different number of slices. If so, we applied a sliding-window approach as illustrated in Fig. 3, left and right panels: The smaller boundary array was compared with all possible chunks of the longer array; the maximum similarity index was finally regarded as the desired measure of similarity.

## Results

### Unsupervised clustering

Figure 4 shows an unsupervised clustering result example for a VISp location, using the simplest neuro-biological feature set F1, that was, only neuron density. The feature values for the different z-slices are shown in panel E of the figure. Panels A and B illustrate the distribution of the clusters and panels C and D the c-layers after the 7^th^ and 8^th^ iteration of the clustering algorithm and the corresponding hierarchical laminar levels, respectively. Panels F and G further represent an overview of the entire iterative clustering process. Specifically, panel F depicts the process in terms of the iteration-specific c-layer distribution, while panel G represented the corresponding global hierarchical tree. Comparing panels F and G to the manual labelling result (shown between panel F and G), it can be seen that our hierarchical tree and especially levels 7 and 8 were in accordance with Brodmann’s classification scheme. Yet, corresponding B-layers merged at different hierarchical levels of our tree. For instance, B-layers IV and Va merged at the second last level, whereas B-layers Vb and VI merged at the last level. In addition, the hierarchical tree also indicated the existence of sub-layers that eventually merged with different 7^th^ or 8^th^ level c-layers compared to B-layers. Figure S1 contains, for instance, a thin c-layer before the 7^th^ level, which, according to manual labelling, should belong to B-layers II/III, but merged with more superficial c-layers at the 7^th^ level. These examples illustrate the potential of the proposed clustering approach to allow deeper insight into the layer structure and its hierarchical organization when compared to the classical approach of manual labelling.

The influence of the feature sets on the clustering and the similarity of the resulting hierarchies are summarized in Fig. 6, using a VISp area data set. It can be seen that feature sets that shared features resulted in more similar trees (Figs. 6A and 6C). Further, regarding κ_m_, the feature sets F1, F2, F3 and F4 yielded much more similar results compared to the manual labelling than feature sets F5 or F6 (p<0.001, t-test with Bonferroni correction); see Figs. 6B and 6D. Compared to F1, the similarity to the manual labelling was, however, only slightly increased for F2, F3, or F4, with the differences not being significant (p>0.2); see Fig. 6D. This, in turn, illustrate that already neuron density alone, among all features investigated, appears to contain most of the information that guided the manual labelling (see also the also the last *Results* section).

### Supervised learning and layer transfer

Similar to the unsupervised clustering results, assignment of d-layers was sensitive to the selection of the applied feature set. In addition, the d-layers depended on the selected reference data set and the reference c-layers, respectively. Yet, the example shown in Fig. 5 illustrated that, if appropriate, neuro-biologically meaningful feature sets were used, the assigned d-layers were similar to the manual labelling results. In the specific case, the VISp data set of Fig. 4 served as reference data set; the respective reference c-layers were to be transferred to a RSC location data set. The unsupervised learning results (using feature set F1) for the RSC data set are shown in panel B of Fig. 5, and the supervised learning results (using feature set F3; results shown for all 200 SVMs) in panel A. The final d-layers were presented in panel D and well agreed with the B-layers indicated to the right of the d-layers.

**Figure 5.**
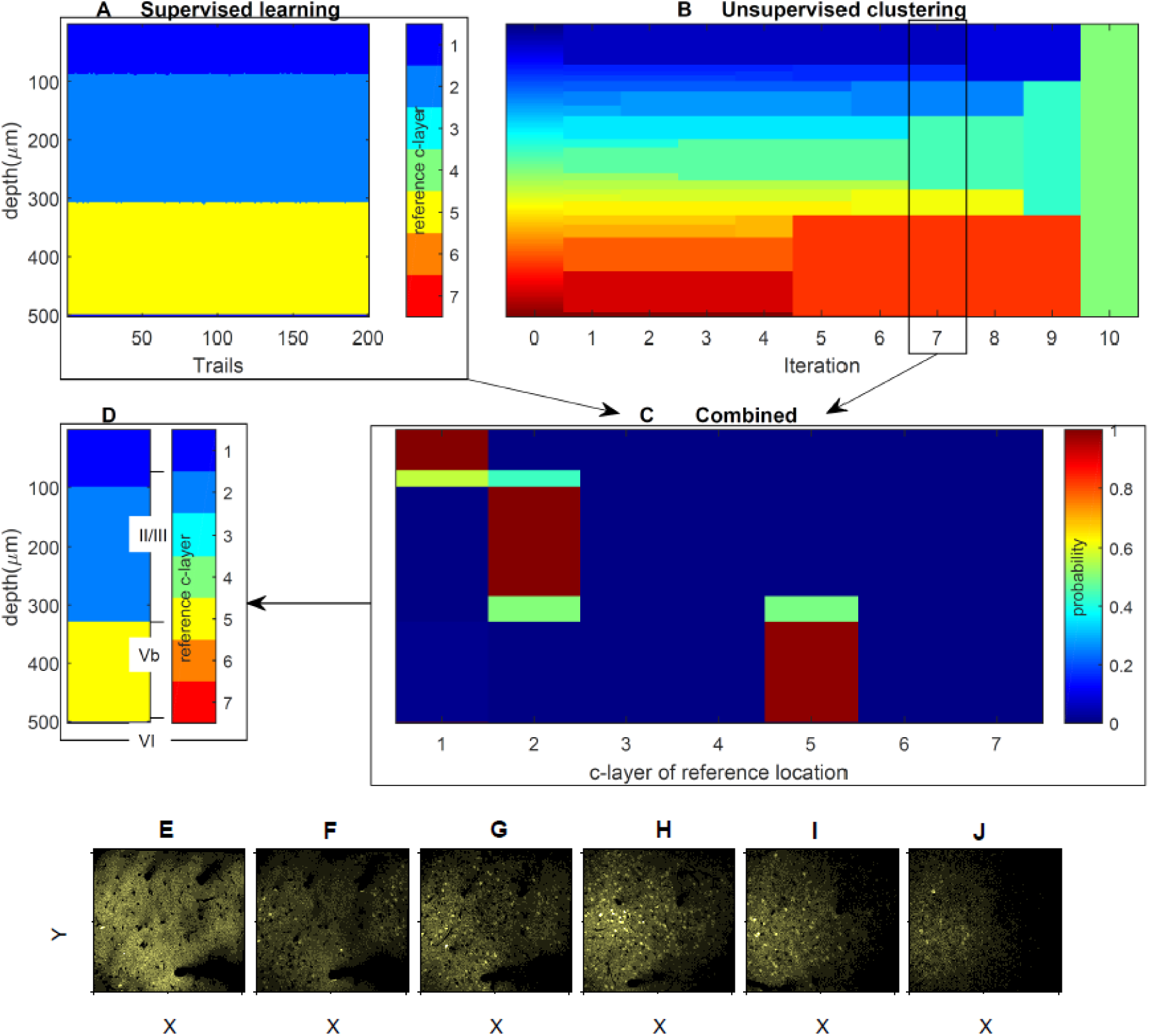
Example for transferring reference c-layers to a target data set. Panels A and B represent the results of the supervised and unsupervised learning of the target data set. Panels C and D show the probability of the assignment of the reference c-layers to the target c-layers as determined by an SVM ensemble and the final discrimination result. Unsupervised clustering was based on feature set F1, and the supervised learning on F3. The dataset under discrimination is the RSC area M262L0, and the reference data for supervised learning are the c-layers of the 8^th^ hierarchical level of VISp area M248L1, that is, the one shown in Figure 2. For comparison purposes, the manually labelled layers are indicated to the right of panel D. The bottom panels show six slice examples at z=50 μm (E), z=90μm (F), z=140μm (G), z=220μm (H) z=300μm (I), and z=400μm (J), corresponding to the six c-layers on the seventh hierarchical level in panel B. Both panels E and F correspond to the first d-layer in panel D, whereas panels G, H, I correspond to the second d-layer and panel J corresponds to the third d-layer.

Different to unsupervised clustering, in the supervised learning part, the single feature *neuron density* was not sufficient to ensure high similarity between d-layers and manual labels. This finding was reflected in the evaluation of the quality measures *n_B_*, *f_c_*_=B_, 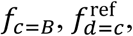, and *f_d_*_=B_, with the results summarized in Figs. 7 (VISp reference data sets) and S2 (SSp reference data sets). In particular, panels A to H in Fig. 7 and panels A to G in Fig. S2 represented the feature set-specific performance in terms of the index *f_d_*_=B_, i.e. the fraction of slices with agreement of the d-layers and the manually assigned B-layers, for the individual VISp and SSp reference data sets. It could be seen that F3 and F4 resulted in higher *f_d=B_* values than F1 for all reference data sets, with the differences being significant (p<0.01) for, e.g., six of the eight VISp reference data sets. However, F4 did not result in a significantly better performance than F3 (p>0.4 in all eight cases), even though F4 included the mean size of all neurons as additional feature. Similar observations held also true for 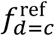.

### Neuro-biological vs. image texture features

Figs. 6, 7 and S2 further reveal that the neuro-biologically inspired features and features sets F1 to F4 clearly outperformed the direct image-based texture features and feature sets F5 and F6 in terms of κ_m_ and *f_d_*_=_*_B_*. Thus, every feature and feature set leaded to a hierarchical tree, but the results were not necessarily neuro-biologically meaningful and comparable to classical layer discrimination patterns.

**Figure 6.**
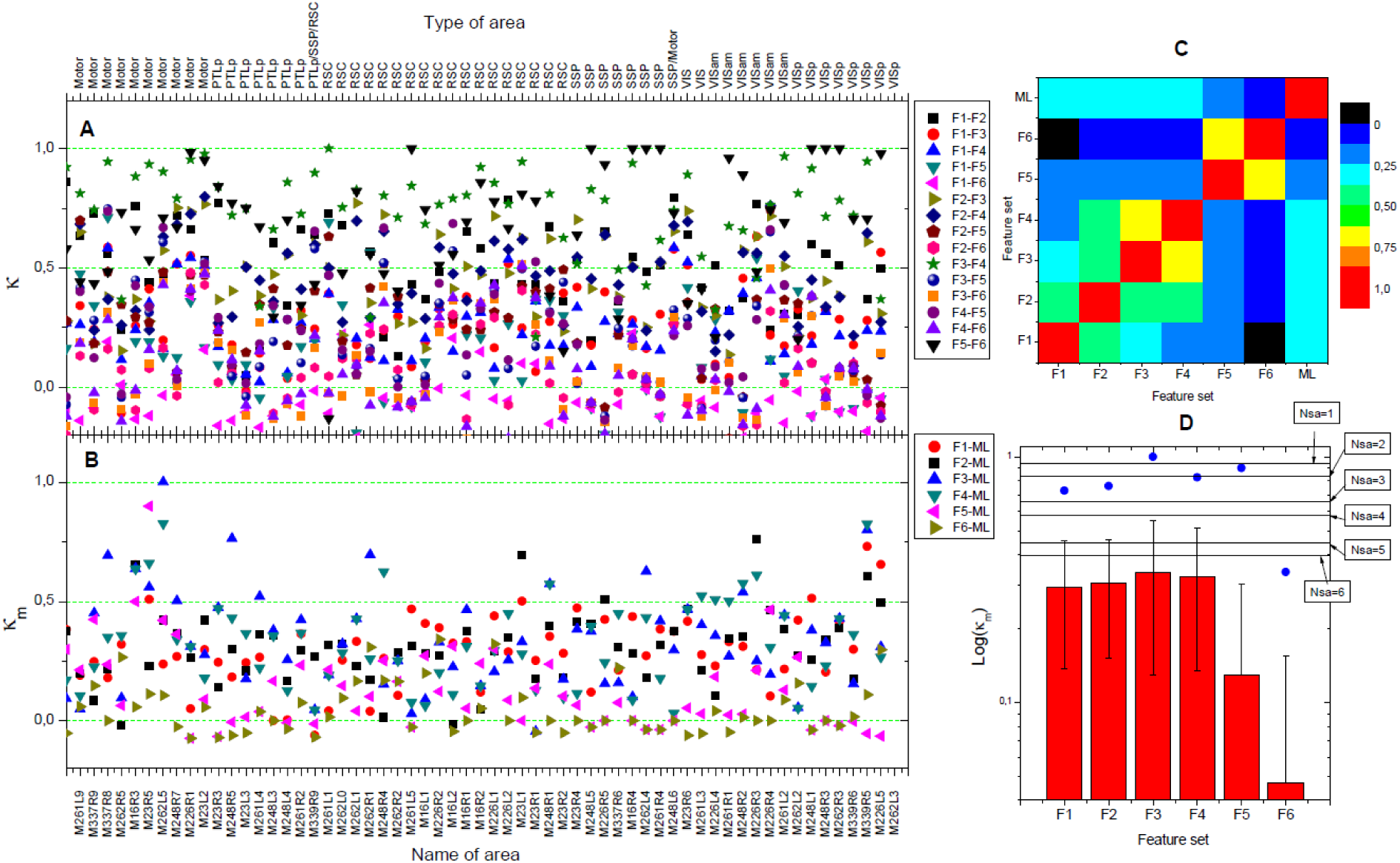
Evaluation of unsupervised clustering. (A) The similarity κ of the results obtained for any pair of feature sets and all 60 data sets studied, and (B) the similarity κ_m_ of the feature set-specific results and the manually labelled layers (indicated with ML). The bottom label of panel B shows the data set names and the top label of panel A their area types. (C) The first six columns and first six rows show the mean values of κ over all 60 locations, and the seventh column and the seven rows the mean value of κ_m_ over all 60 locations. The element at (ML, ML) is manually set to 1. (D) The detailed statistics of Log(κ_m_). The histograms show the mean values, error bars the standard deviations, and the blue dots the maximum values. The minimal values are close to zero or even negative, and are, therefore, not shown. Pairwise comparison of F1, F2, F3, and F4 values is not significant (p>0.2), whereas pairwise comparison of F1, F2, F3, or F4 to F5 or F6 is significant (p<0.001). To illustrate the meaning of the obtained κ_m_ values, we randomly shifted the manually assigned layer boundaries, with the boundary shift values chosen in the range of [-Nsa, Nsa] slices. We repeated runs with Nsa = 1,2, …, 6. For each Nsa value, each of the 60 locations was re-labelled and the artificial labelling result compared to the original manual labelling. Corresponding mean values are plotted in panel (D). Based on our experience, a NSA value of 5 approximately resembles the accuracy of manual layer discrimination; the respective line therefore represents ideal performance when taking into account manual labelling uncertainty.

**Figure 7.**
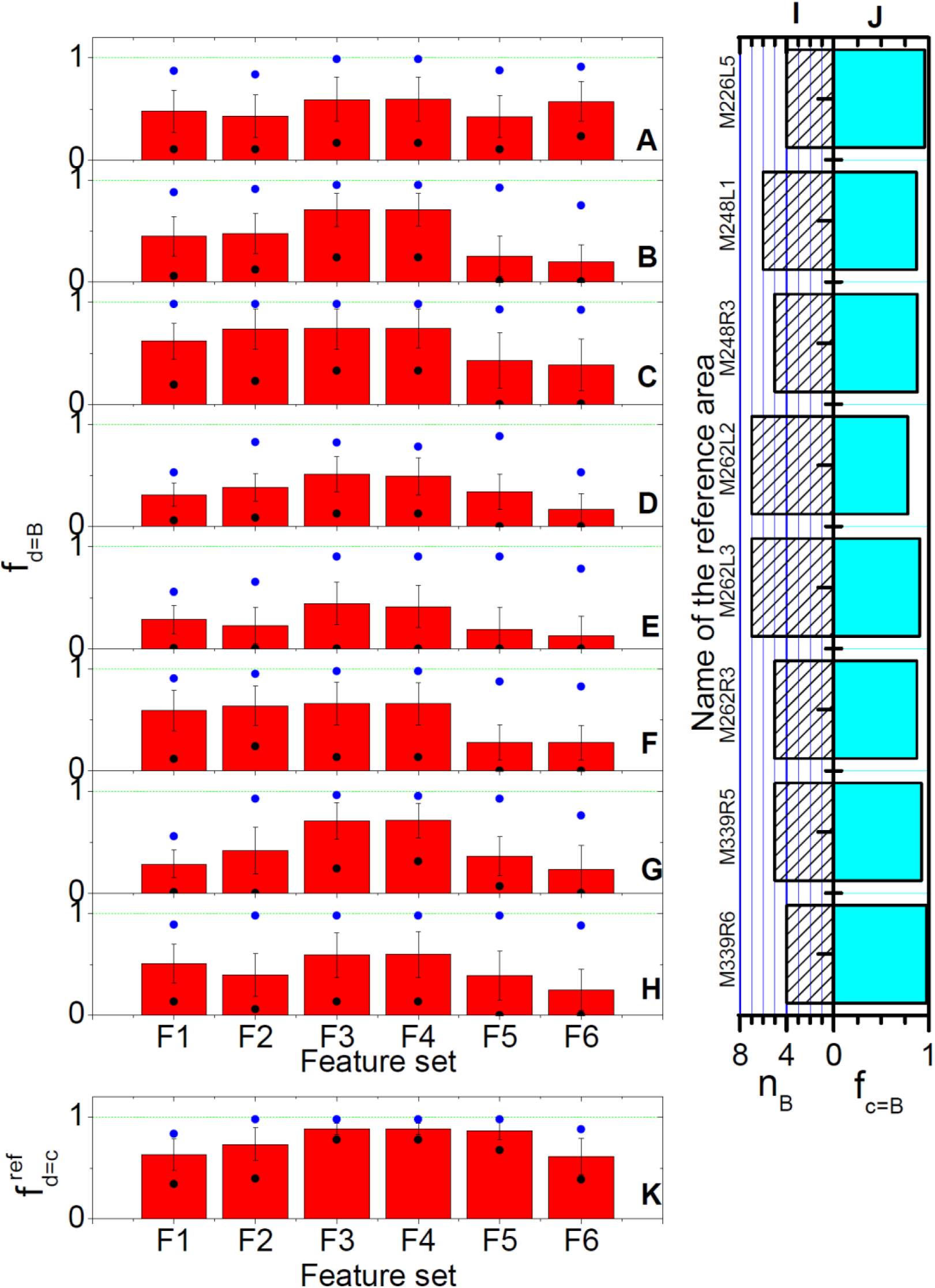
Supervised learning evaluation results for the VISp reference data and c-layers. (A-H) *f_d=B_* statistics for the different reference locations (panels) and different feature sets (horizontal axis). The histograms show the mean values over the 60 locations, the error bar the standard deviations, and blue and black dots the maximum and minimum values. Comparison of differences between F1 and F3 or F1 and F4 are, except for M262R3 and M339R6, significant (p<0.01). In contrast, differences between F3 and F4 are not significant for all cases (p>0.4). Panels (I) and (J) contain additional information about the reference c-layers. Panel (I) shows the number of manually discriminated and labelled layers of the reference locations, namely n_B_. (J) shows the similarity between the refined reference c-Layer structures and the refined manually labelled layers structures. (K) The statistics of the self-identical measurement (= re-labelling) of the reference locations, where all the symbols have the same statistical meaning as in panels (A-H). All reference c-layers were selected on the hierarchical level after the 8^th^ clustering iteration. Target c-Layers were selected on the last hierarchical level that exhibited a layer number not smaller than the number of reference c-Layers.

This observation is further supported by Fig. 8, which represents cross-location similarity of d-layer-based discrimination results. Here, feature set F1 was used for clustering, F3 for supervised learning and an SSp area was chosen as reference data set. The resulting pattern revealed high similarity between image data from locations of the same area type, and thereby resembled the expected results and the corresponding pattern obtained by the manual labels (see supplemental Fig. S3), although corresponding observations after unsupervised clustering and hierarchy comparison were not that obvious (Fig. S4). Comparison of Fig. 8 and Fig. S5 further demonstrate that the results were also sensitive to the selection of the reference c-layers, but, more importantly, application of the direct image-based features did not lead to a (neuro-biologically) meaningful pattern at all (Fig. S6).

**Figure 8.**
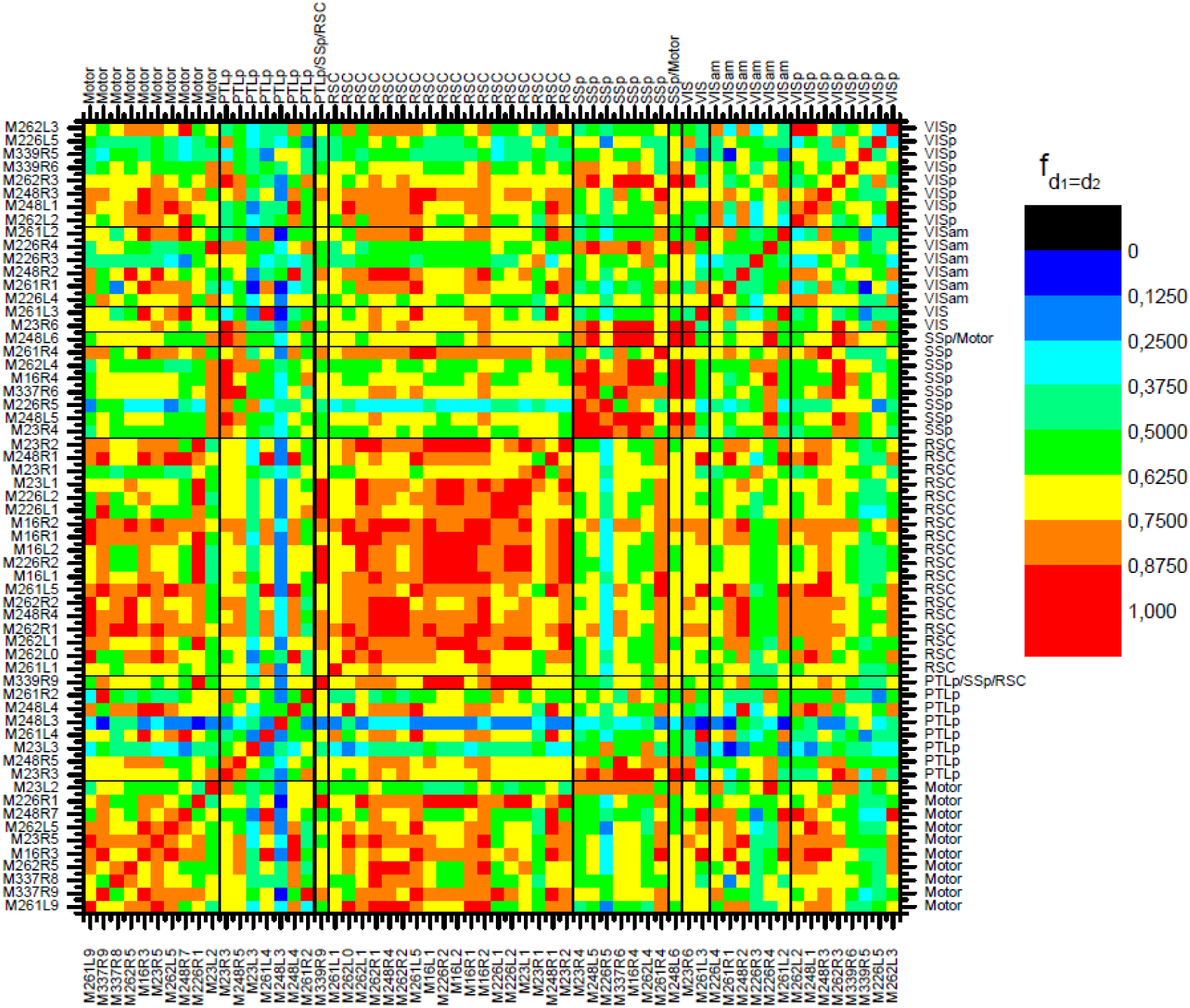
Similarity of discrimination results for any pair of datasets, using feature set F3 for supervised learning and M16R4 as reference data set. Reference c-Layers were selected after the 8^th^ clustering iteration, and all target c-Layers were selected on the last hierarchical level with the number of layers not being smaller than the number of the reference c-Layers.

## Discussion

Machine learning methods that are able to discriminate cortical laminar patterns and their hierarchy are of high demand in the field of neuroscience^38-41^. In this study, we demonstrate that combining unsupervised clustering and supervised learning is well suited for such tasks. The main ideas are that iterated unsupervised clustering allows appropriately describing the laminar hierarchical structure for individual cortical locations, and supervised learning employing “rich structure” reference locations enables to bridge those hierarchical patterns between different locations. Last but not least, our results also illustrate that selecting neurobiologically meaningful feature sets is an important prerequisite for obtaining neurobiologically meaningful layer discrimination results. Subsequently, some more specific advantages as well as interesting aspects and potential shortcomings of our approach are discussed in more detail.

### No need for manual labelling

One significant advantage of our layer discrimination procedure is that there is no need to manually identify and label the layers. Therefore, the method is less limited by subjective perception and related potential observer bias. Simultaneously, the procedure provides extended information on the hierarchy of laminar cortical organization and opens up more flexible ways to study cortical structure-function relations. Naturally, human interaction is still required to select the hierarchical level and reference locations and layers, depending on the particular research interest, but this is inevitable and not considered as a shortcoming of the method itself.

### Slice-versus (individual) neuron-based layer discrimination

In this work, we use vertical image slices as objects of interest to perform the discrimination task, which is different from alternative approaches that detect and then assign individual neurons to different layers^40^. In comparison, assigning neurons to layers usually requires quite a number of elaborated features, whereas in this work we eventually used five or fewer features (F1 has one feature; F3 has five features) to obtain promising results. In addition, the number of neurons is naturally larger than that of the image slices; in our case, the number of neurons in a single image data set is on the scale of ten thousand, whereas the data set comprises only a few hundred slices. Therefore, advantages of using slices include (depending on the complexity of considered neuron features), for example less computation time and simpler feature selection. In contrast, a shortcoming is that in the horizontal direction (i.e. within slices), the entire slices are assumed to be homogeneous regions that can be unambiguously assigned to a specific layer. Alternatively, smaller regions or patches of the slices with more homogeneous characteristics could be selected for clustering.

A related aspect refers to a more philosophical issue. As is well known, there exist transition zones between the cortical layers, which cannot unambiguously be assigned to specific layers. During neuron-based laminar discrimination, transition layers could in principle be defined as regions where upper and lower layer neurons are entangled together. Nonetheless, such compartments would usually be assigned to either the upper or the lower layer, based on some implementation-specific criteria^40^. In our approach, transition layers automatically show up as separate layers at more detailed hierarchical levels, but may merge with either the upper or the lower neighbouring layer at higher hierarchical levels. As an example, in Figs. 4, 5 and S1, such layers can be easily identified.

### Impact of feature selection: What means “meaningful” layer discrimination?

It needs to be noted that, especially for the unsupervised clustering approach, it is not easy to evaluate the appropriateness of a feature set. Any (combination of) features will generate a hierarchical tree. In our case, we evaluate the appropriateness of features mainly by their ability to generate a final single-level layer discrimination that resembles manually labelled (Brodmann) layers. Human observers (and most probably also Brodmann himself one century ago)^48,49^ discriminate cortical layers mainly visually based on neuron density and neuronal morphology. Thus, it was expected that corresponding features performed best in terms of respective quality measures. Yet, other features and feature sets do not necessarily have to be less meaningful, but can potentially reveal additional, currently neglected structuring principles. Finally, to judge appropriateness and importance of feature sets and discrimination results, it remains to be seen and investigated whether there exists, for example, a functional correspondence of the uncovered hierarchy and layer discrimination.

Still, based on similarity to manual labelling results according to Brodmann’s classical laminar classification scheme, we tentatively conclude that F1 (neural density), F2 (neural density and mean neuron size), F3 (neural density, the proportions of the two neuron classes, and the mean neuron sizes of the two classes), and F4 (F3 plus the mean size of all neurons) give promising results, and, if computation cost is considered critical, neuron density alone already leads to reliable laminar hierarchies by the proposed unsupervised clustering approach. For the described supervised learning part, we, nevertheless, demonstrate that F3 and F4 significantly improve performance compared to neuron density alone. In contrast, the image texture features and F6 do not lead to satisfying results at all.

Considering that our applied methods of neuron size and shape detection and neuron classification are not very precise at the moment (as the development of such algorithms is not our main goal and beyond the scope of this work), we can reasonably expect that developing better methods will further improve the presented discrimination results.

## Conclusion

Combining unsupervised and supervised machine learning allows discriminating (and finally analysing) the hierarchical laminar cortical organization in 2-photon image data without the need of manual labelling. Unsupervised clustering offers the opportunity to identify the hierarchical laminar organization of image data obtained at the different cortical locations. Regarding respective features for layer discrimination, it turned out that neuron density alone was already sufficient to achieve biologically plausible results during unsupervised clustering, whereas supervised learning for cross-location label transferring requires consideration of additional features. The computed discrimination results agree well with the classical Brodmann scheme of cortical layers and reflect the layer similarity of data acquired at locations of the same type of cortical area. At the same time, they also provide additional, new information on the laminar structure of cortical areas, for instance, by providing insight into the hierarchical structure revealed by the clustering analysis.

## Acknowledgments

This work is funded by German Research Foundation (DFG) and the National Natural Science Foundation of China in the project Cross-modal Learning, DFG TRR-169 / NSFC (61621136008) - A2 to CCH and JSG, DFG SPP2041 as well as HBP/SP2 SGA2, DFG SFB - 936 - A1, Z3 to CCH, NSFC (31671104) to JSG, and DFG SFB -1328 - A2 to RW. The authors thank Changsong Zhou for helpful discussions and Farid I. Kandil for his suggestions on the denotation of the different types of layers.

## Author contributions

J.-S. G. and C. C. H. designed the research; D. L., M. Z., and R. W. developed the discrimination methodology; G. W., Y. H., and H. X. worked on the experiments and pre-processed the image data; D. L., R. W., J.-S. G., and C. C. H. wrote the paper; all authors approved the paper.

## Supplementary Information

**Table S1.** Refined maps between reference c-Layers from SSp areas and the manually labelled layers. Similar to the VISp areas, some layers were merged after visual inspection.

| Reference location name | Reference c-Layer structure | Manually labelled layer structure |
| --- | --- | --- |
| M23R4 | [1], [2], [3], [4, 5] | [1], [2], [3], [5] |
| M248L5 | [1], [2], [3], [4, 5] | [1], [2], [3], [5] |
| M226R5 | [1, 2, 3], [4], [5] | [1, 2, 3], [4], [5] |
| M337R6 | [1], [2], [3], [4], [5] | [1], [2], [3], [4], [5] |
| M16R4 | [1], [2], [3], [4], [5] | [1], [2], [3], [4], [5] |
| M262L4 | [1], [2], [3], [4, 5] | [1], [2], [3], [5] |
| M261R4 | [1], [2], [3], [4], [5] | [1], [2], [3], [4], [5] |

**Figure S1.**
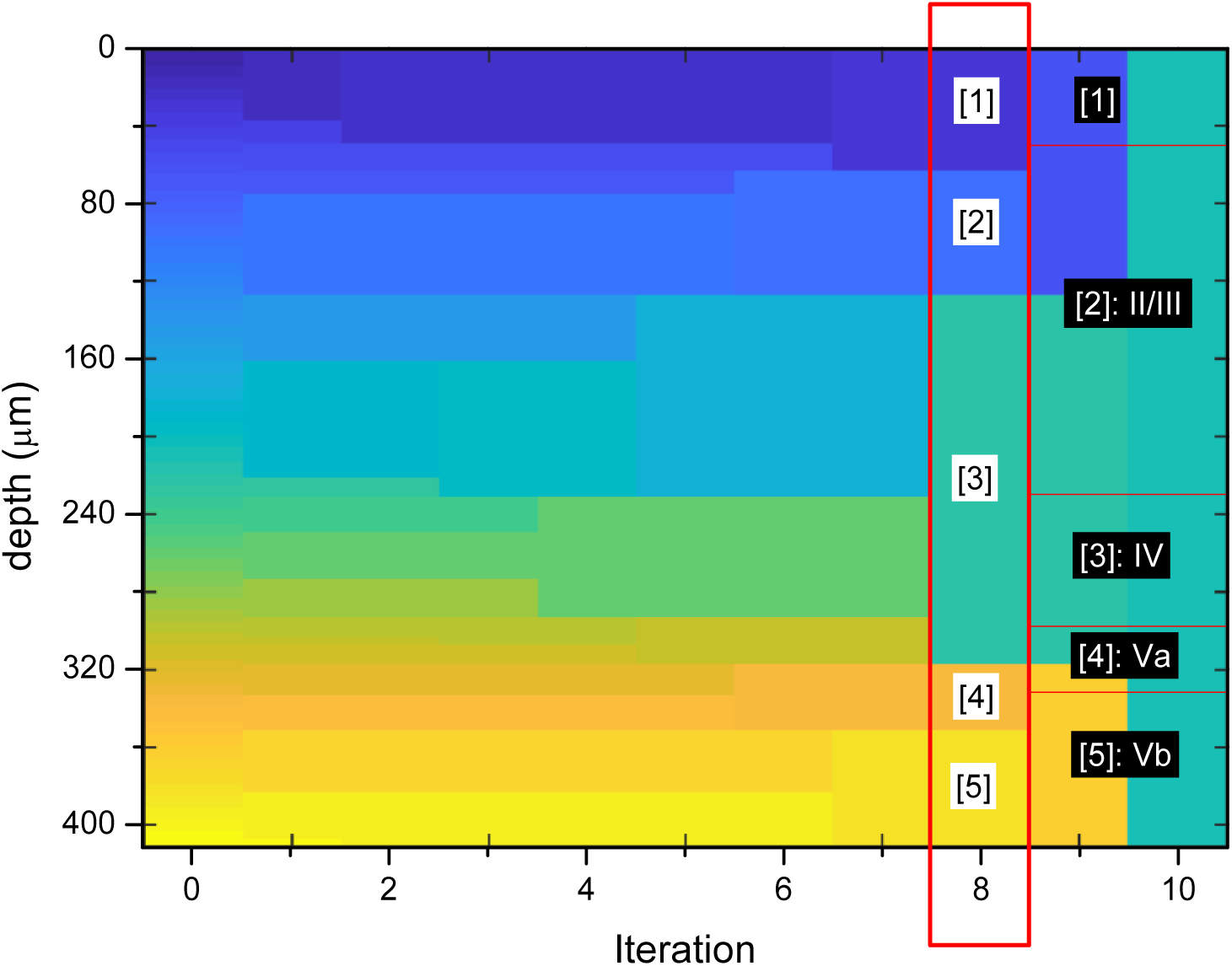
Comparison between unsupervised clustering results of M248R3 with feature set F1 and its corresponding manually labelled layers. The red rectangle highlights the reference c-layers we used (layer names indicated by black text with white background). To its right side, the red horizontal lines indicate the B-Layer boundaries (layer names by white text and black background).

**Figure S2.**
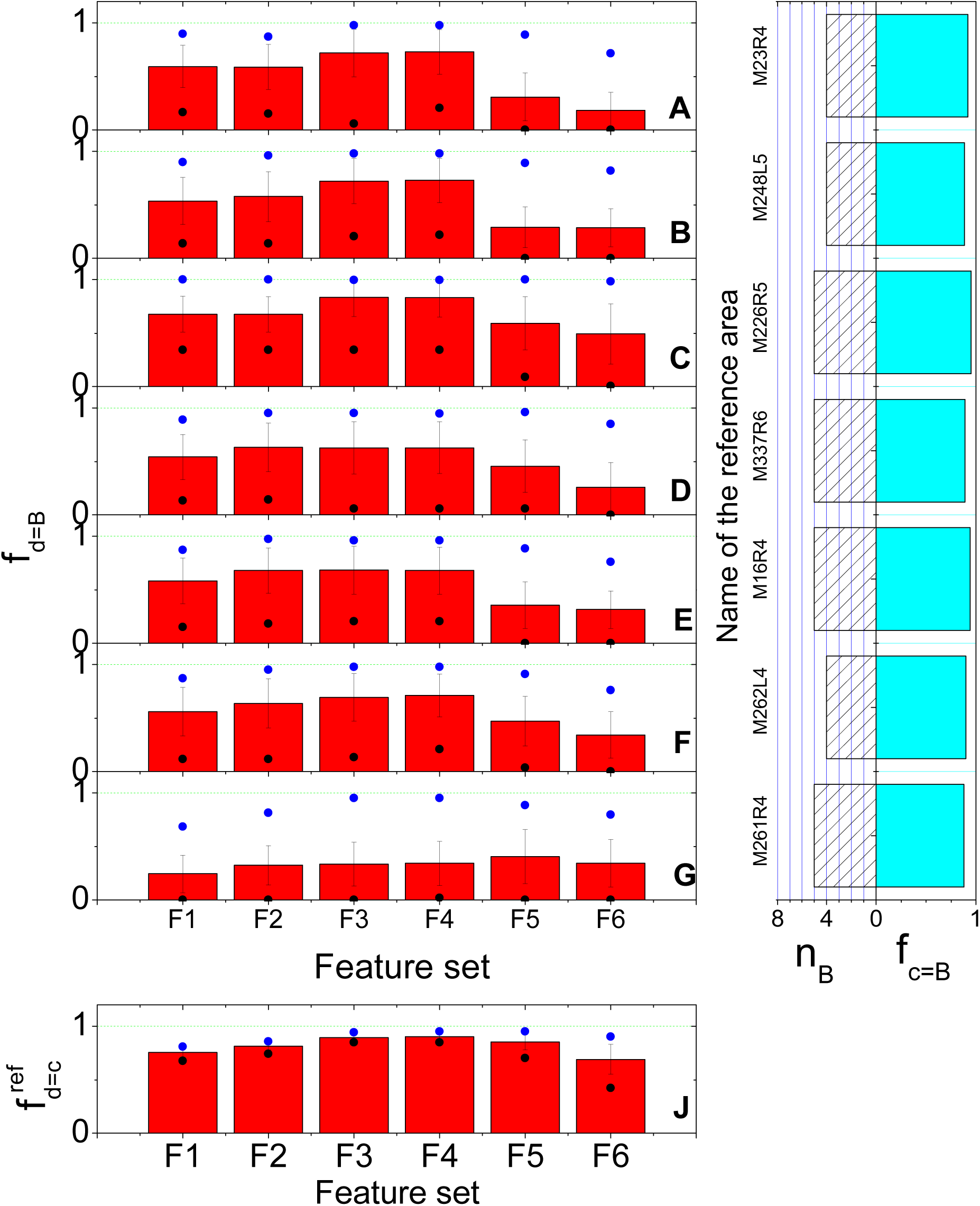
Supervised learning part evaluations for SSp reference c-Layers. All figure details and symbol meanings similar to Figure 5.

**Figure S3.**
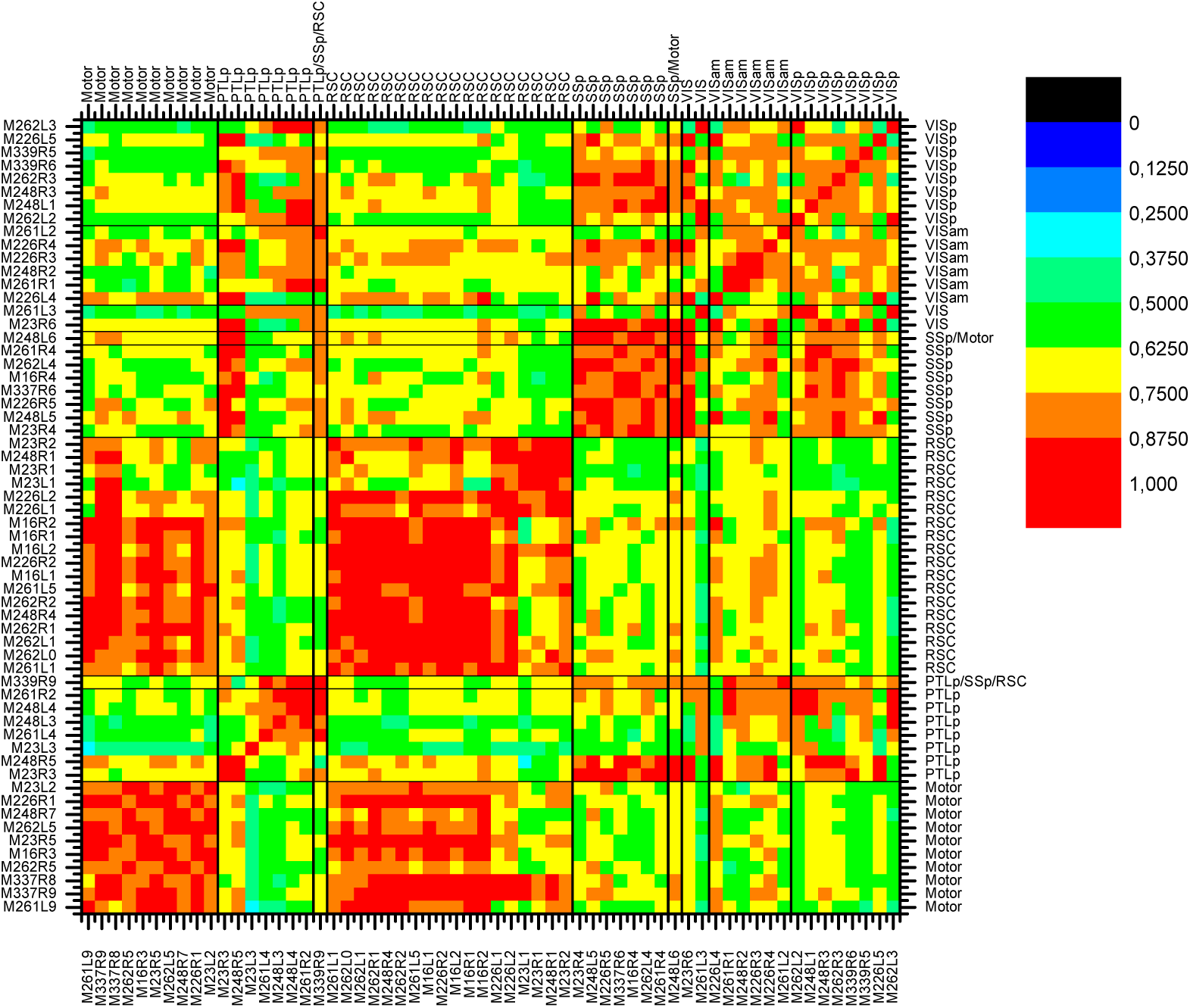
Similarity of manual labels for any pair of locations.

**Figure S4.**
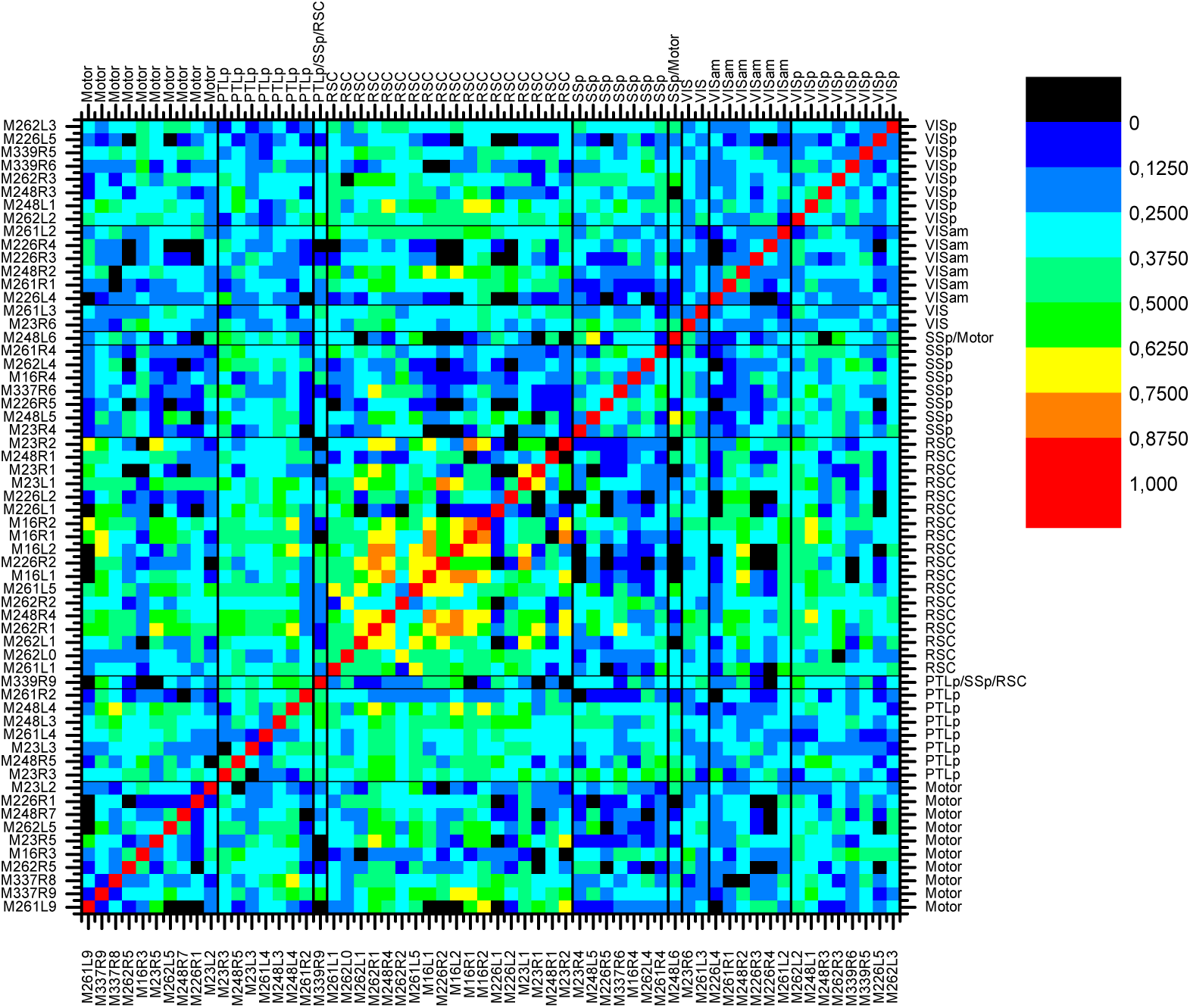
Similarity hierarchical clustering results for any pair of datasets and using feature set F1 for clustering.

**Figure S5.**
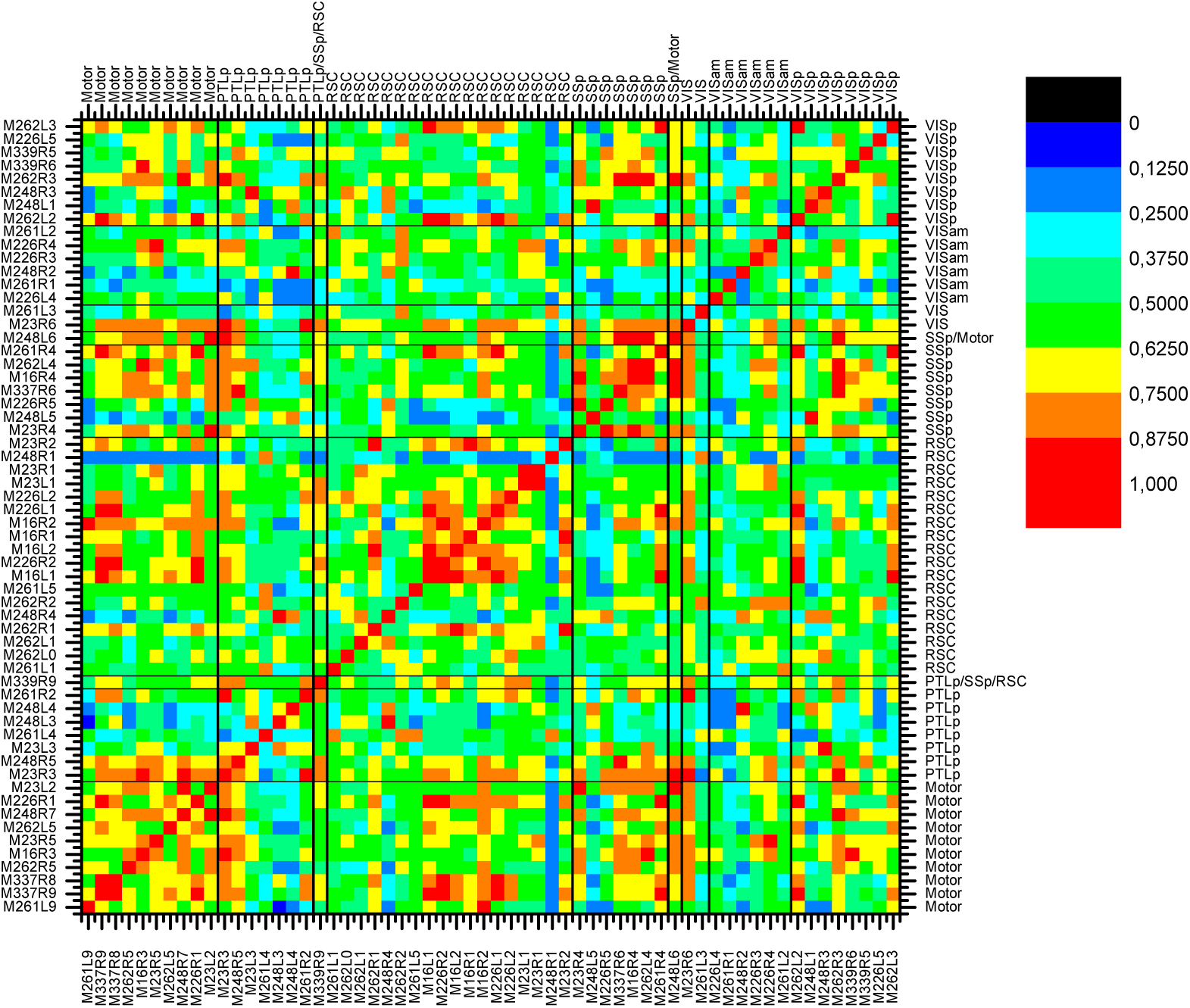
Similarity of layer discrimination results for any pair of dataset locations, using feature set F3 for supervised learning and taking M226R3 as reference location.

**Figure S6.**
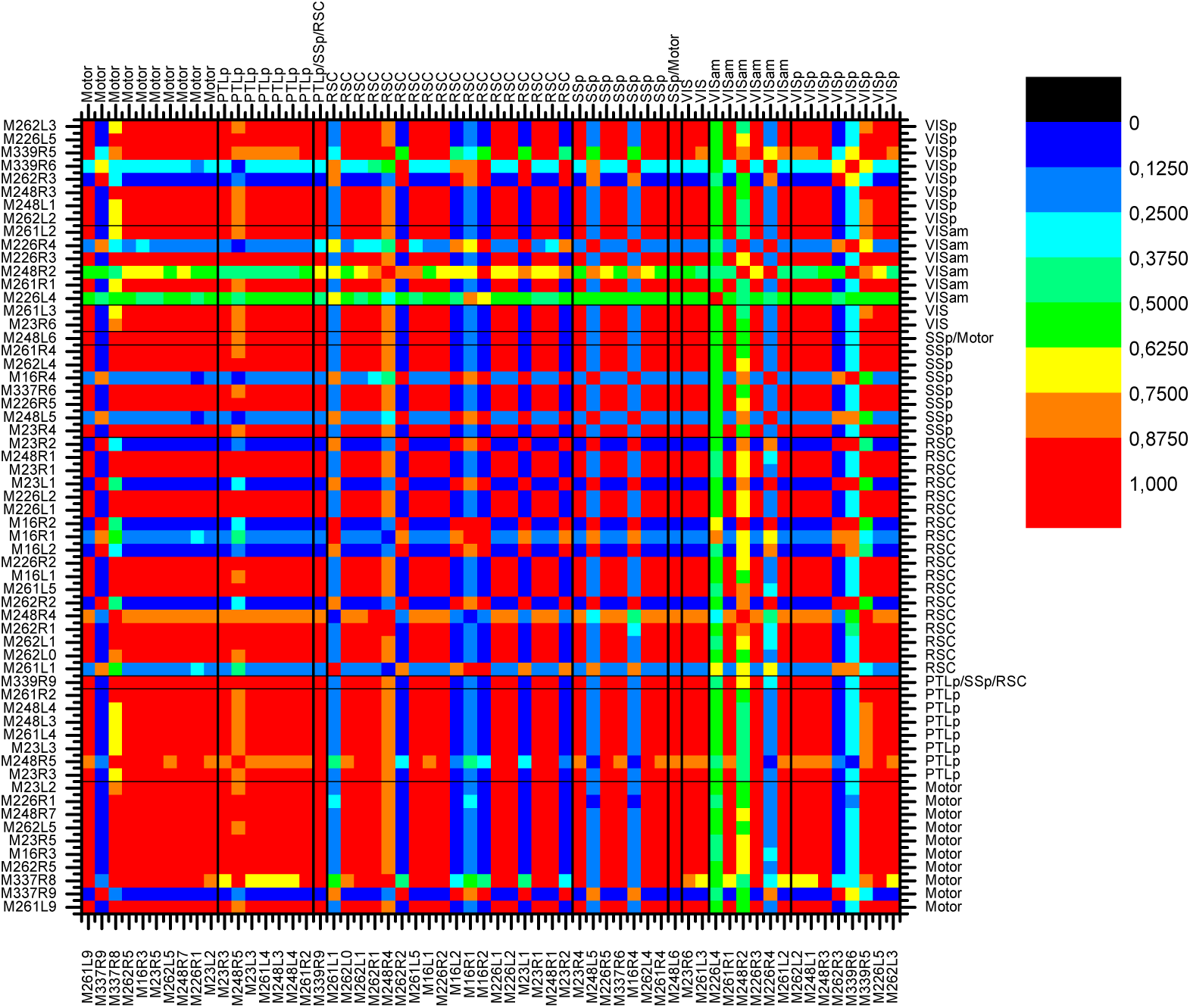
Similarity of discrimination results for any pair of dataset locations, using feature set F6 for supervised learning and taking M16R4 as reference location.

